# Assembly of minimally fragmented metagenomic pangenome graphs

**DOI:** 10.64898/2026.10.29.762972

**Authors:** Dylan Duchen, Florencia Martino, Kakhangchung Panmei, Abraham J. Kandathil, Steven J. Clipman

**Author notes:** **Corresponding Author:** Dylan Duchen, PhD, MPH 300 George Street, #505 New Haven, CT 05611, USA.

## Abstract

Pangenome variation graphs are a powerful tool for representing genetic diversity within and across populations. For comparative and evolutionary genomic analyses leveraging these graph-based structures, it is important that graph topology reflects the underlying phylogenetic and genetic similarities of the genomes embedded within the graph. However, in metagenomic settings these approaches often produce highly fragmented multi-component graphs, hindering comparisons of genomes embedded across different components. This challenge is particularly pronounced for hypervariable viral families like *Anelloviridae*, whose short circular genomes frustrate existing variation graph construction methods. We present PANIMaSTA (Progressive Average Nucleotide Identity and Maximum Spanning Tree-Based Assembly), a workflow to construct minimally fragmented metagenomic pangenome sequence variation graphs through progressive assembly. The order in which the genomes are incorporated into the graph is informed by a maximum spanning tree (MST) inferred from an all-to-all genome similarity matrix comprised of pairwise *Anelloviridae* average nucleotide identities (ANI). Applied to over 5,200 diverse *Anelloviridae* genomes, we construct a sequence variation graph whose graph topology reflects the global genome similarities observed across the ANI matrix. PANIMaSTA constructs minimally fragmented metagenomic pangenome sequence variation graphs that are not achievable with existing methods suitable for similarity-aware sequence-to-graph mapping and downstream genetic analyses across highly diverse genomes.

## Introduction

Pangenome variation graphs have emerged as a transformative approach for capturing genetic diversity across populations, enabling comprehensive representation of both major and minor genomic variations within a single non-linear structure.(*1–3*) Unlike traditional linear reference genome representations, pangenome graphs integrate multiple genomes into a unified joint sequence space, facilitating analyses of genetic variation, evolutionary relationships, and population structure. This approach is particularly critical for metagenomic datasets, where high levels of genomic diversity challenge conventional alignment-based methods.(*4–7*)

The utility of pangenome graphs is particularly pronounced for hypervariable viral families, such as *Anelloviridae*. Members of this viral family have short circular DNA genomes, ranging from 1.5 to 4 kb, and exhibit high levels of sequence diversity.(*8*) Despite their ubiquitous presence in mammalian hosts, the evolutionary and ecological dynamics of *Anelloviridae* remain poorly understood.(*9*, *10*) Existing classification guidelines, such as those from the International Committee on Taxonomy of Viruses (ICTV), rely on pairwise nucleotide identity thresholds to delineate species boundaries, with a 69% identity cutoff frequently used for open reading frame 1 (ORF1).(*11–13*) However, novel Anelloviridae species continue to be discovered, underscoring their vast genetic diversity and the limitations of existing alignment-based methods to perform intra- or inter-species analyses.

Current state-of-the-art tools, such as PGGB and Minigraph-Cactus have demonstrated utility for constructing pangenome graphs from closely related genome collections;(*14*, *15*) however, their reliance on all-versus-all pairwise alignment followed by graph induction (PGGB) or on incremental alignment to a reference-seeded backbone (Minigraph-Cactus), was designed to resolve variation across similar genomes and does not extend to metagenomic settings, where many highly divergent, species-representative genomes are embedded in a single graph. When genomic divergence spans wide evolutionary distances, these approaches produce highly fragmented multi-component graphs, reducing their utility for downstream applications such as metagenomic read alignment, taxonomic classification, and evolutionary inference.

These limitations are especially acute for the small, circular genomes of the *Anelloviridae*, one of the most genetically diverse viral families known, whose diversity is further shaped by extensive recombination.(*9*, *10*, *12*). While Progressive Cactus offers a viable approach to sequence variation graph construction for highly diverse genomes, it uses a phylogenetic tree to infer intermediate ancestral genomes which are embedded into the graph and the order by which these genomes are incorporated.(*16*) However, this assumes a guide tree can be reliably constructed and the ancestral states accurately inferred across deep divergences, both unlikely for hypervariable viral families. Ordering inputs to guide progressive graph construction is not itself new: Virus-VG greedily ordered contigs by maximal overlap before progressive construction with VG msga to avoid disconnected components,(*17*) and PanGraph progressively merges sequence subgraphs along a similarity-derived guide tree without reconstructing ancestral genomes.(*18*) These address within-host quasispecies assembly and bacterial pangenomes, respectively, rather than a family-wide reference spanning genus-level divergence.

To overcome these challenges, we developed PANIMaSTA (Progressive Average Nucleotide Identity and Maximum Spanning Tree-Based Assembly). This workflow integrates 1) pairwise all-to-all global average nucleotide identity (ANI) estimation across the sequence set using VCLUST, (*19*) 2) clustering and the identification of representative genomes using MMseqs2, (*20*) 3) the construction of a ANI weighted graph/network, 4) maximum spanning tree (MST) inference, 5) a frontier expansion approach to traverse the MST and identify the order these genomes are progressively assembled into a sequence variation graph, and finally 6) sequence-to-graph mapping and embedding of the non-representative genomes. PANIMaSTA infers a trajectory through the ANI-based MST to maximize the relatedness of each genome incorporated into the graph compared to the existing set of embedded genomes, resulting in a minimally fragmented sequence variation graph. In doing so, PANIMaSTA avoids the need for any ancestral-genome reconstruction while producing a graph whose topology reflects the global genome-to-genome similarities observed across the full metagenomic space.

As a stringent test case, we applied PANIMaSTA to 5,217 *Anelloviridae* genomes, demonstrating its ability to generate a single component sequence variation graph that captures the full breadth of genomic diversity, evaluating performance in terms of graph connectivity and fragmentation relative to the distribution of ANI similarities observed across the database. We show PANIMaSTA provides a minimally fragmented graph whose topology reflects the ANI similarity matrix. We show that graph-space distance between embedded genomes is inversely correlated with their pairwise ANI, that this relationship is retained for sequences aligned to the graph after its construction, demonstrating the resulting structure is calibrated to genome relatedness. Our findings highlight the utility of PANIMaSTA for constructing unified pangenome variation graphs, with broad implications for metagenomic studies involving highly diverse viral or microbial populations.

## Methods

### Dataset

*Anelloviridae* genomes comprising this dataset (N=5,217) reflect complete circular genomes of 1.5 to 4 kb infecting multiple vertebrate species. The majority of the *Anelloviridae* genomes were sourced from the SCANellome database (Anellovirus_2024.2, N=3,864),(*21*) with the remainder being obtained from the National Center for Biotechnology Information (NCBI) nucleotide repository using the accession IDs listed in previous publications or the ICTV9 reference set.(*9*, *11*, *12*) Each genome underwent initial quality control to remove duplicated genome sequences using seqkit.(*22*)

### PANIMaSTA Workflow

Briefly, pairwise average nucleotide identity (ANI) estimation is performed on all genome pairs using VCLUST.(*19*) All-to-all ANI estimation is performed over all genomes to obtain a complete pairwise similarity matrix. Clustering is performed to identify a representative subset of genomes to avoid over-representing any one species during core pangenome graph construction. The similarity matrix, limited to the representative genomes, is then used to construct a weighted undirected graph using genomes as nodes and pairwise ANI values as edge weights using the iGraph R package (v2.3.1). The edge weights are then rank transformed and a maximum spanning tree (MST) is derived from the weighted graph, whereby the remaining edges reflect the highest pairwise ANI observed between genomes. From the representative genome set, the core pangenome graph can be constructed by selecting an initial seed genome and iteratively incorporating its neighbors in the MST using a frontier-expansion approach to embed the most similar genome available. While building the full graph is an option, partitioning the MST into genus-like subtrees, building each subtree in parallel, and then embedding genomes across the subtree-specific graphs is the default pangenome assembly approach. In this approach, edges below a pairwise ANI similarity are cut, resulting in subtrees solely comprised of edges above the threshold. The resultant structure is used to inform subtree-specific graph assemblies. Genomes lying at the edges of each subtree-subtree interface, along with any isolated genomes with no edges above the ANI threshold, are split into chunks and aligned against its neighboring subtree graphs and then stitched together to form a bridge between the subtree graphs. After core graph construction, the non-representative genomes are incorporated into the graph through sequence-to-graph alignment to both obtain the complete pangenome graph and confirm its topology corresponds to the MST network and global pairwise ANI similarity matrix structures.

### Sequence preparation and representative genome inference

Input genomes are first passed through a preparation step that normalizes sequence identifiers and removes redundancy. Sequences are deduplicated by content and non-alphanumeric characters in the FASTA sequence headers/identifiers are replaced. Every record is renamed under the PanSN specification (sample#haplotype#contig) so that path names remain unambiguous through graph construction, alignment, and downstream indexing. A mapping table between original and sanitized identifiers is retained so that results can be traced back to source records. All 5,217 Anellovirus genomes were carried forward for pairwise all-to-all ANI estimation using VCLUST to derive a matrix of all genome-to-genome total ANI (TANI) estimates. Because VCLUST estimates TANI by Lempel-Ziv-based alignment (LZ-ANI) rather than k-mer sketching, the identity estimate does not depend on a chosen k-mer size or iterative k tuning, a property well suited to short circular genomes for which sketching approaches are sensitive to the choice of k; k-mer-based similarity estimation via kmer-db2 is used as an initial step in VCLUST to prefilter the provided database.

Representative genomes are inferred by clustering the prepared sequences with MMseqs2 at a default minimum sequence identity of 70%, just above the ORF1-specific 69% pairwise identity threshold used by the ICTV to delineate Anelloviridae species, so that each species-level lineage includes a distinct representative. For this dataset, 5,217 input genomes were reduced to 1,749 representatives carried forward for graph construction. Cluster assignment is also written to a table used during downstream analysis to assess the concordance between sequence-to-graph alignment-based genome embedding and the initial MST topology.

### Network construction, subtree partitioning, and tip folding

The MST inferred over the ranked TANI network can be partitioned into subtrees by removing edges whose TANI falls below a pairwise similarity threshold. The threshold may be supplied directly or derived from the pairwise TANI distribution. TANI threshold derivation involves masking uninformative pairwise similarity estimates (TANI <0.25), as TANI values below this range suggests any successful alignments between these pairs of genomes would reflect spurious relatedness.

After low-TANI masking, density antimodes are searched to locate gaps between modes of the distribution. Additionally, by iteratively pruning the MST edges across a range of thresholds, an exact component curve can be obtained. Any plateauing in the number of obtained components across the TANI search space reflects TANI estimates over which the partitioning does not significantly alter the number of components. Relatedly, we can assess the effect each TANI pruning threshold has on the size of the largest subtree/component in order to identify the ranges resulting in a similar plateauing of primary component size. This primary component size analysis, termed percolation plateau identification, focuses on the stability of the graph components rather than simply the total number of components. Percolation-based threshold selection is the default approach for determining the TANI threshold, since a threshold associated with large subtree stability is highly informative for identifying genus-level thresholds.

For the *Anelloviridae* database, the procedure returns a genus-level value of 0.397, supported by a percolation plateau spanning 0.370 to 0.460 within which the largest component holds at 27.3% of genomes and estimates delineating genus-level differences in the literature.(11) (**Supplementary Figure S1**) Removing edges with TANI ≤0.397 produced 114 subtrees, of which 40 were carried forward for progressive graph assembly. The number of assembled subtrees is below the total number of components because pruning MST edges can result in isolated tips or islands, so any subtrees with ≤3 genomes are folded into the nearest subtree with the highest TANI edge. Folded-in genomes and genomes incident to these removed edges are designated ‘bridge’ genomes, and the specific subtree pairs each bridge genome connects are recorded. Because a single genome may be incident to several removed edges, bridge genomes are prioritized by the number of distinct subtree pairs they span, and a degree-aware set-cover selection identifies a minimal subset sufficient to cover every cross-subtree edge in the partitioned MST. Folded-in and edge-defined bridge genomes are split into chunks and each chunk mapped to the adjacent component graphs they connect to, so that no representative genome is excluded from core graph construction. The MST, its partitioning, and the assembly order within each component are rendered as layout plots for inspection, with nodes colored either by component membership or by position in the within-component assembly order (**Figure 1**).

**Figure 1.**
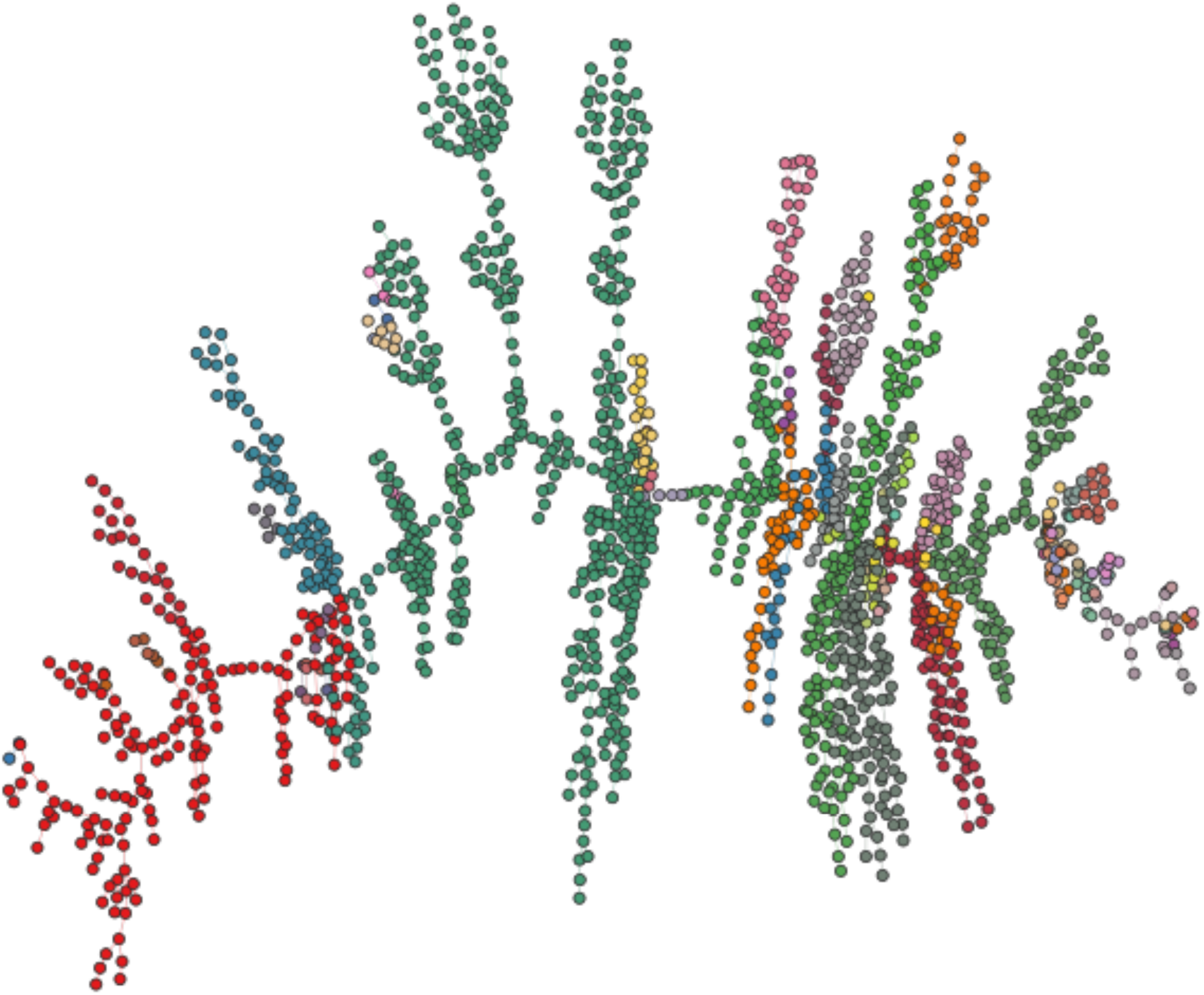
Topology of the PANIMaSTA MST graph. Maximum spanning tree layout of the 1,749 representative anellovirus genomes, with nodes colored by component membership inferred using a minimum TANI edge threshold of 0.397.

### Graph construction

PANIMaSTA supports two main assembly modes that use MST-derived similarity structure. In ‘--simple’ assembly mode, no partitioning is performed and the full MST is traversed by frontier expansion from a seed representative genome. Progressing through all 1,749 representative genomes, the unincorporated genome with the highest TANI to any genome already embedded in the graph is aligned and incorporated into the core sequence variation graph via VG msga. An initial seed selection step evaluates several candidate starting genomes that have non-overlapping early orderings to retain the candidate whose early expansion maximizes the mean TANI identity to the already-embedded set. This is performed since the ordering of the early set of genomes determines the initial node space into which all subsequent genomes are aligned (**Supplementary Figure S2**).

In the subtree-partitioned assembly modes ‘--isolated’ and ‘--mst_assembly’, each component is assembled independently in parallel. Within each component, the seed is the genome with the highest mean incident-edge TANI, and the remaining genomes are ordered by the same greedy frontier expansion albeit restricted within the set of genomes associated with the subtree. Component FASTA files are written in this order and passed to VG msga. If the VG msga construction yields a multi-component graph for a specific subtree, the largest component is isolated and the genomes associated with any smaller components are incorporated into the primary component via sequence-to-graph mapping and augmentation using VG.

While the ‘--isolated’ mode combines the resultant component graphs into a single sequence variation graph file comprised of a unified node ID space and no edges between components, ‘--mst_assembly’ re-establishes the cross-subtree connections defined by the MST. The bridge genomes are divided into non-overlapping chunks, and these chunks are aligned with VG map against the component graphs corresponding to the subtrees associated with each bridge genome. Chunked alignments are filtered by score, and a connectivity check verifies that each required cross-subtree edge is covered before augmentation proceeds. Where a selected bridge genome fails this check, an alternative bridge genome covering the same edge is substituted if available. Component graphs are then augmented with their assigned chunked alignments, embedding bridge-derived sequence into the node space of adjacent components. Node identifiers are re-unified across the augmented components and the graphs are combined, at which point the chunked paths belonging to each bridge genome are stitched into a single contiguous path so the complete genome is embedded. Because these stitched paths traverse node space contributed by two different components, they contain the edges that render the combined graph traversable across subtree boundaries.

Bridge genomes whose chunked alignments were not selected or successfully aligned to the components of interest are incorporated into the graph via multiple rescue approaches.

Rescue is first attempted by sequence-to-graph alignment and augmentation by extracting a subgraph comprised of the genomes assigned to the same subtree/component and using this for sequence-to-graph alignment. The extracted subgraph retains the global node and edge IDs, enabling the rescued genome to be incorporated into the full graph despite only being mapped to the relevant component-specific genomes. As component-targeted rescue requires at least one of the genomes assigned to the component to be embedded in the full graph to serve as an anchor, if all the genomes associated with the component are absent then rescue is attempted through sequence-to-graph alignment against the whole graph. All sequence-to-graph alignment uses X-drop extension and a band-width longer than the largest genome being aligned.

A circularization option is provided by the user to make every path in the representative and complete graphs a closed circular walk. As all anellovirus genomes are circular, we used this option to circularize each genome incorporated into the graph during assembly and after augmentation by adding an edge joining each path’s terminal node to its first. Aside from assembly, an explicit circularization step is carried out on the completed representative graph and after all remaining genomes are embedded.

### Embedding remaining genomes

Two graphs are produced by PANIMaSTA. An initial graph comprised of the MMseqs2-representative set (N=1,749) and a graph comprised of the complete deduplicated genome set. For the Anellovirus database, the 3,468 non-representative sequences are embedded by sequence-to-graph alignment using VG map and augmentation. All 5,217 genomes are embedded into a unified pan-*Anelloviridae* graph this way, and the resulting paths circularized as above, yielding a final graph in which the complete dataset is represented while the topology is determined by MST-guided pairwise global ANI similarities.

### Completeness and path naming

Two properties of the delivered graph are enforced explicitly rather than assumed. Completeness is verified by comparing the path set against the input FASTA, and any genome still absent is aligned and embedded in a final pass, so the graph contains every input genome exactly once. Duplicate path names are removed, keeping one copy where the walks are identical and reporting the conflict where they differ, since the indexer skips rather than rejects duplicates and a graph can otherwise report the expected path count while loading with genomes missing. Finally, path/genome naming is harmonized. Internally, the pipeline assigns short temporary IDs for genomes used in progressive assembly to avoid potential name-parsing errors, while genomes added by alignment retain their PanSN names and augmentation appends a subrange suffix. All output graphs use the PanSN-formatted names of the input FASTA.

### Graph assessment

Assembled graphs were evaluated on three criteria. First, component structure and path inventory were recorded to quantify the degree of fragmentation, taken as the number of connected components, and to verify that each genome is represented by a single path, with path names reconciled against the input FASTA so that absent or spurious paths are reported explicitly. Second, embedding of the non-representative genomes was assessed by the proportion that aligned to the representative graph, by the fraction of each genome’s own length covered by its alignment, and by the distribution of alignment identity across thresholds, so that the assessment reflects how many sequences were correctly placed and at what sequence-to-graph similarity threshold. Third, topological concordance was quantified by computing the mean nearest-node distance (MNND) in traversal steps through the graph for pairs of embedded genomes and correlating it against the pairwise TANI for the same genome pairs.

Pairs were sampled across TANI strata, 25 per 1% TANI bin within each pair class, so that the full range of divergence is represented rather than the high- or low-identity comparisons that might dominate the pairwise space, and MNND was estimated over five bootstrap replicates, each sampling 25% of the nodes on a genome’s path, with a floor of 20. A further set of pairs was added so that every aligned genome has at least one representative partner, taken as its five nearest by TANI. These guarantee coverage for the placement analysis and are excluded from the correlation estimates. Pairs were reduced to a single orientation before any correlation, and 95% confidence intervals were obtained by the Fisher z transform on the resulting count of unique pairs. Pairs in separate isolated components with no finite traversal distance (i.e. when PANIMaSTA is run in ‘--isolated’ mode) are excluded from the correlations. Alongside MNND, a neighbourhood overlap estimate was computed across several hop radii using overlap estimates of the node sets traversed by each pair (Jaccard and Dice). Jaccard comparisons were partitioned on whether the compared set of genomes fell within the subtree, as the shared node space would therefore track relatedness. Concordance between each genome’s nearest neighbors by TANI (as defined by the MST) and by graph-space proximity (as defined by the assembled sequence variation graph) is also computed, with correlations reported overall and stratified by whether both, one, or neither genome in a pair belonged to the representative set.

### Alternative graph construction approaches

PANIMaSTA was compared with PGGB, Minigraph-Cactus, and Progressive Cactus. PGGB constructs graphs from all-to-all genome-wide alignments, retaining only mappings above a user-specified identity threshold. Minigraph-Cactus is reference-anchored, incrementally aligning each assembly to a graph seeded by a designated reference genome and inserting the divergent segments that align, with base-level variation added by Cactus in a subsequent step. Progressive Cactus instead aligns genomes along a guide tree, reconstructing ancestral sequences at internal nodes to span divergence that cannot be aligned directly. For Progressive Cactus we used a guide tree inferred for the 1,749 representative *Anelloviridae* genomes using ROADIES.(*23*) Resulting graphs were evaluated on component structure and on the number of input genomes successfully embedded.

## Results

### Single-Component Pangenome Graph Construction

In simple assembly mode, PANIMaSTA generated a single-component pangenome graph from the 1,749 representative anellovirus genomes, into which all 5,217 genomes in the dataset could subsequently be aligned. Despite the extreme diversity within this viral family, the maximum spanning tree approach bridged highly divergent nodes through intermediate sequences, producing a unified graph whose topology recapitulated the global genetic and phylogenetic similarity structure spanning the breadth of anellovirus diversity. Visualization of the final structure revealed tight clusters corresponding to closely related species, as well as longer paths that linked more distantly related clades. Because the maximum spanning tree retains every genome as a node and connects each through its highest-identity available neighbor, no pruning of the MST was required, no genome is lost during network construction, and no tip-removal criterion or component folding is applied. Component-specific assembly reached the same endpoint, yielding a single connected graph after bridge-mediated stitching and only 2/1749 genomes requiring remapping before embedding.

### Global Topology Retention with Component-Specific Assembly

Because the simple assembly mode threads all clades through a single spanning backbone, the most divergent genomes are necessarily incorporated into one common graph, which can place genomes with little shared sequence in close graph-space proximity. This is avoided by the ‘--mst_assembly’ approach, where the partitioning threshold was derived from the TANI matrix data. A percolation-focused approach over the pairwise TANI estimate across the representative genomes identified a plateau from 0.370 to 0.460 across which the size of the largest connected component changed by less than 2% per 0.01 TANI. Refinement within that window by MST edge-gap analysis selected 0.397 (threshold values from 0.395 to 0.411 provide comparable partitions). Cutting the 1,748-edge MST at this threshold removed 113 edges and yielded 114 subtrees, of which the largest contained 477 genomes. The 1,635 retained edges carry TANI values concentrated around 0.6. Across all pairwise comparisons rather than MST edges alone, within-subtree pairs had a median TANI of 0.239 (n=155,653) against 0.037 between subtrees (n=1,372,973, AUC 0.976). The within-subtree median sits below the threshold because subtrees are connected components, so only 7.5% of within-subtree pairs are themselves above it while the rest are joined through chains of intermediates. As expected when cutting a maximum spanning tree (equivalent to single-linkage clustering at the threshold), none of the 11,696 pairs at or above the threshold crossed a subtree boundary, so no directly similar pair was separated by the cut (**Supplementary Figure S1**).

Of the 114 subtrees, 40 held enough non-bridge genomes to be assembled as independent graphs, covering 1,554 genomes, while the remaining 74 had their members chunk-mapped into adjacent components instead, 23 holding two or fewer non-bridge genomes and 51 composed entirely of bridge genomes. Component assembly, bridge mapping and stitching together completed in 58 minutes, since each VG msga invocation was presented only with genomes spanning within-genus divergences rather than the full breadth of the family.

Removing the partitioning edges defined 47 component-to-component relationships the final graph was required to represent, and degree-aware set-cover selection over 97 candidate bridge genomes identified 42 sufficient to cover all of them. Chunk-level alignment of these against the component graphs they connect, followed by augmentation and path stitching, re-established connectivity across subtree boundaries while embedding each bridge genome as a single contiguous path, incorporating all 1,749 representatives into one component. Unlike simple assembly, the partition is cut at 0.397, genome pairs below that threshold are connected only through bridge genomes rather than through shared node space, preserving separation between divergent genera while retaining high-resolution structure within them (**Figure 1**).

### Embedding of the Full Dataset by Sequence-to-Graph Alignment

Graph construction uses only the 1,749 representative genomes, so a practical requirement is that the remaining 3,468 non-representative sequences can be placed into that node space without degrading it. Aligning these sequences against the assembled graph with VG map embedded all 3,468 as new paths, yielding a final graph containing all 5,217 genomes in the dataset. Every sequence aligned with no unmapped genomes. Alignment identity was at least 0.70 for 99.8% of sequences, at least 0.80 for 93.4%, and at least 0.90 for 64.1%. Each aligned genome traversed existing nodes where it shares sequence with an embedded representative and contributes new nodes where it diverges, so the node space of the final graph remains anchored to the representative set while the path set spans the complete input. All paths, including those introduced by alignment, were circularized, so every genome in the final graph is represented as a closed circular walk consistent with the biology of the family.

That non-representative genomes aligned successfully is expected given that each was assigned to its representative at 70% identity during clustering, but the relevant observation is where they align. For 98% of mapped genomes, the nearest representative in graph space belonged to the same MST-derived subtree as the representative they were assigned to during clustering, and the representative itself ranked among the five nearest in 96.8% of cases. The exact assigned representative was the single nearest core path for only 34.6% of mapped genomes, which is expected where a genus contains many representatives, so graph proximity resolves reliably to the subtree rather than any singular genome within it. Node sharing was correspondingly specific, with 22.4% of within-subtree pairs sharing any node space at a median TANI of 0.60 against 0.6% of cross-subtree pairs at a median TANI of 0.11. Mapped genomes therefore recruit node space already traversed by members of their own subtree, indicating that the representative graph captures the sequence space of the clusters it stands in for. This is the property that makes the representative-set strategy tractable, since the progressive alignment cost scales with the 1,749 representatives, while the resulting graph remains a valid alignment target for the full dataset and, by extension, for sequences absent from the current database.

### Graph Topology Recapitulates Pairwise Sequence Similarity

A pangenome graph is only useful as a similarity-aware reference if proximity within the graph carries information about sequence relatedness. We therefore quantified the correspondence between graph-space distance and sequence identity directly, computing for each pair of embedded genomes the mean nearest-node distance (MNND) in traversal steps and comparing it against the pairwise total ANI (TANI) estimated by VCLUST. Pairs were drawn by stratified sampling, 25 per 1% TANI bin within each pair class, so that the full range of divergence is represented rather than the high-identity comparisons that dominate the pairwise space, yielding 7,101 unique pairs. Across the full diversity range spanned by the dataset, MNND was strongly and inversely correlated with TANI (Spearman ρ = −0.589, P < 2×10-16), and median MNND declined monotonically across increasing TANI strata defined relative to the genus threshold. Genome pairs at TANI 0.849 and above were separated by a median of 0.57 traversal steps, pairs at 0.699 to 0.849 by 3.35 steps, pairs at 0.397 to 0.699 by 304 steps, and cross-genus pairs below the 0.397 threshold by 1,038 steps, a 1,806-fold decrease from the lowest to the highest stratum (**Figure 2A**). Genomes that are close in sequence space are thus close in graph space.

**Figure 2.**
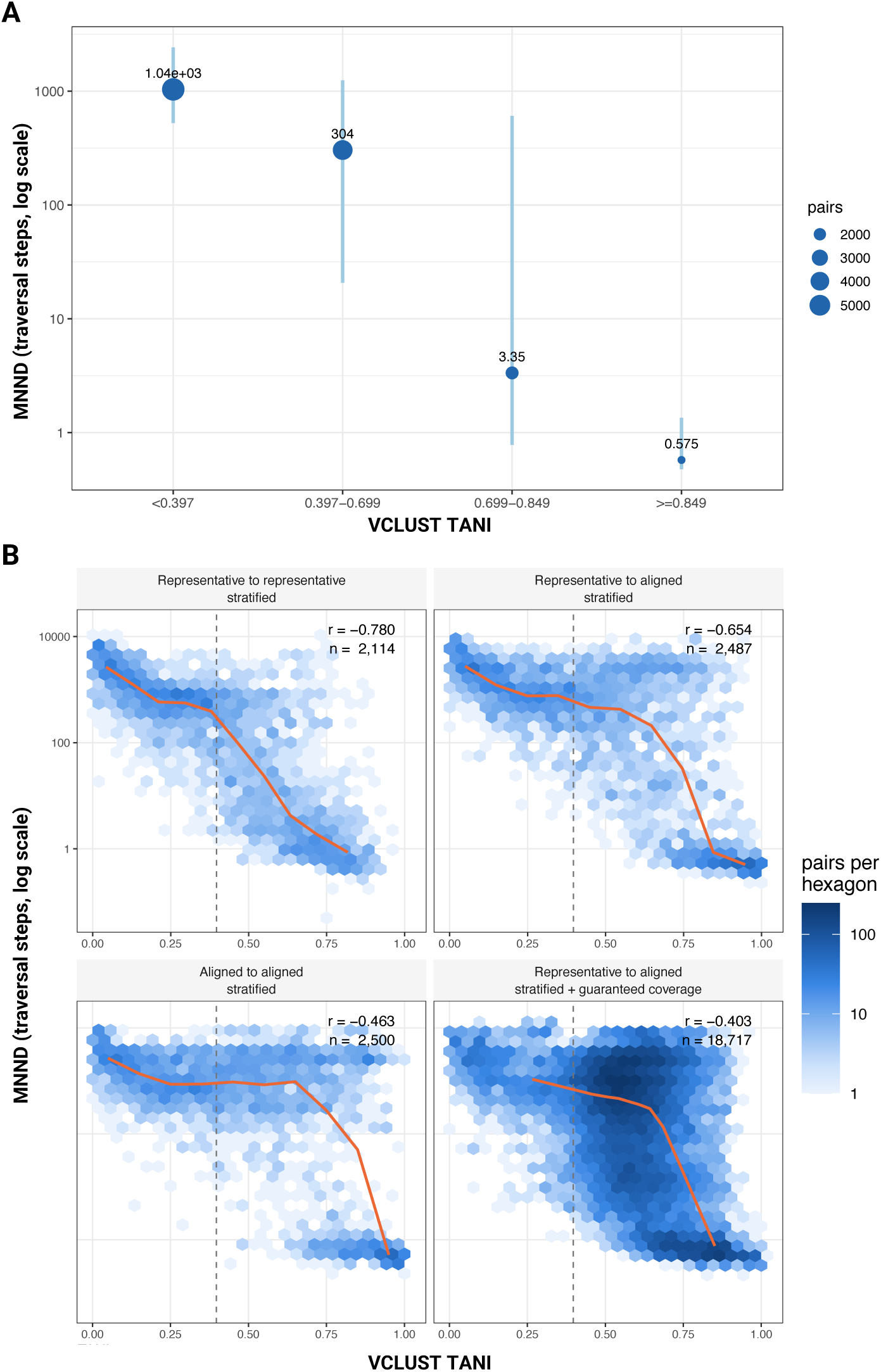
Graph distance reflects pairwise genome similarities. **(A)** Median MNND by TANI stratum with interquartile ranges, log scale. Strata are defined relative to the derived genus threshold of 0.397. **(B)** MNND against pairwise TANI, hexbinned, for pairs stratified by whether both, one, or neither genome belonged to the representative set, with the Spearman correlation and unique pair count for each class. The fourth panel repeats representative-to-aligned pairs with the guaranteed-coverage pairs included; those are each aligned genome’s five nearest representatives by TANI and are range-restricted by construction, which is why their correlation is weaker. One orientation per pair throughout. The dashed line marks the partition threshold.

This relationship held for genome pairs involving sequences that were not used to construct the graph. Stratifying pairs by whether both, one, or neither genome belonged to the representative subset used for progressive assembly, the inverse MNND to TANI relationship was preserved in each stratum (Spearman ρ = −0.780, −0.654 and −0.463 over 2,114, 2,487 and 2,500 unique pairs for representative-to-representative, representative-to-aligned and aligned-to-aligned pairs respectively), with non-overlapping 95% confidence intervals, indicating that the sequence-to-graph alignment step places non-representative genomes at positions consistent with their sequence identity to the embedded representatives rather than at arbitrary locations (**Figure 2B**). Assessed directly, the core genome nearest in graph space to each of the 3,467 aligned genomes belonged to that genome’s assigned subtree in 98% of cases, and its own cluster representative ranked among the five nearest core genomes for 96.8% (**Figure 3**). Read- or genome-level assignment against this graph therefore recovers a position that is interpretable in terms of genome-wide sequence similarity, which is the property required for graph-based classification of novel or divergent anellovirus sequences.

**Figure 3.**
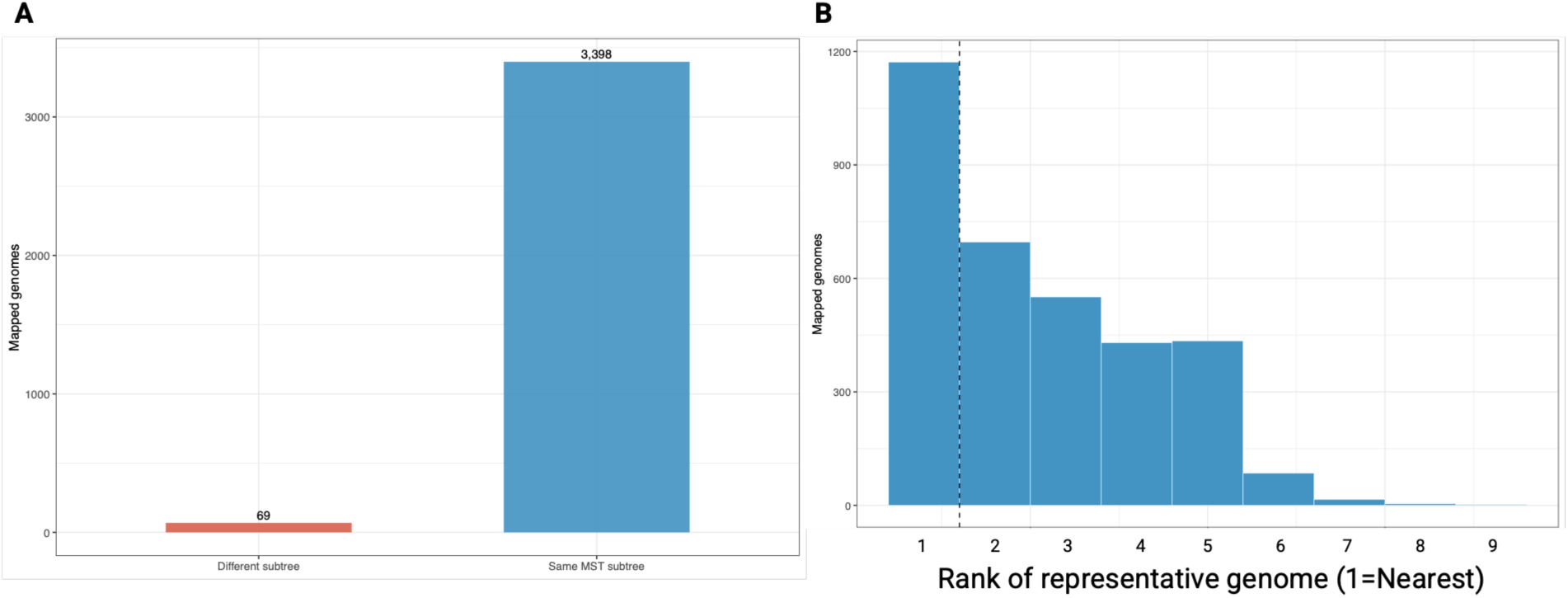
Placement of genomes embedded by sequence-to-graph alignment. **(A)** Counts of aligned genomes by whether their graph-nearest representative falls within an assigned subtree, 3,398 concordant against 69 discordant of 3,467 (98.0%). Bridge genomes count toward both subtrees they connect. **(B)** Rank distribution of each aligned genome’s own cluster representative among those it was compared against, n = 3,391, where rank 1 is nearest in graph space.

### Comparison with alternative graph construction approaches

Applied to the same 1,749 representative genomes, none of the three comparator methods produced a single connected graph spanning the *Anelloviridae* database. PGGB, which relies on all-to-all genome alignments, returned 1,077 discrete components even at the most permissive alignment parameters available (**Figure 4**), 952 of which (88.4%) contained a single genome. Genomes in separate components share no coordinate system, so no traversal, distance, or shared-node comparison can be made between them, and at 1,077 components the majority of the pairwise space in the dataset is inaccessible to graph-based analysis.

**Figure 4.**
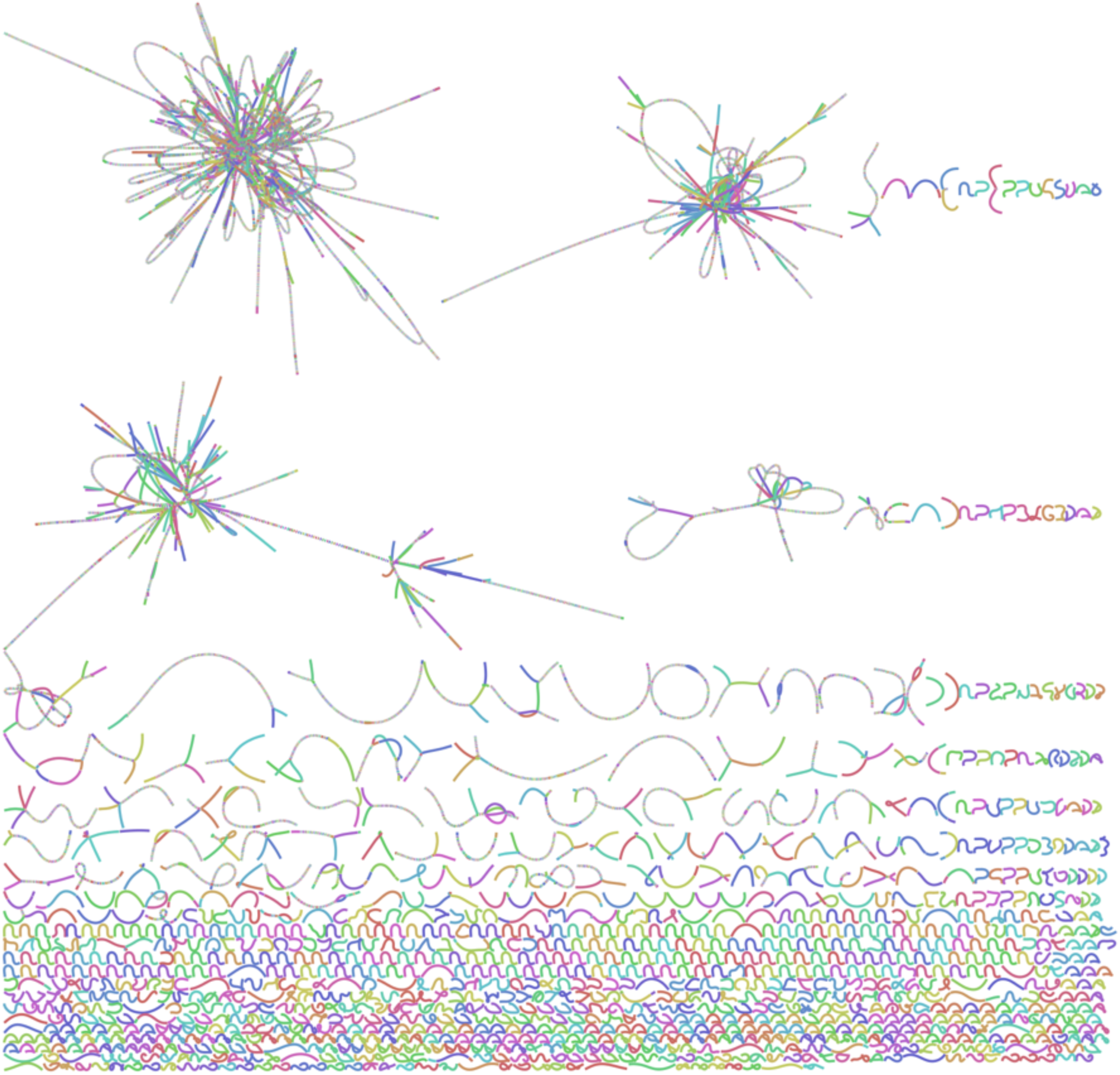
Fragmentation of the PGGB graph over the same 1,749 representative genomes. BandageNG visualization of the PGGB graph comprised of 1,077 discrete components of which 952 (88.4%) contain a single genome.

Minigraph incorporated only those genomes that aligned to the designated reference backbone, so the graph it produced covered a small fraction of the input. Minigraph-Cactus, which builds on that backbone, returned 1,749 independent components, one per input genome. Progressive Cactus completed alignment against a ROADIES-inferred guide tree, but we were unable to convert the resulting Hierarchical Alignment Format (HAL) output into a sequence variation graph.

### Runtime

A direct runtime comparison against these tools would not be informative, since the graphs differ in construction and in what they represent, and a shorter wall-clock time would not indicate that the same work was performed more efficiently. However, whether the workflow completes within a time acceptable for its intended use is informative. For the anellovirus database, the full pipeline from sequence preparation through embedding of the non-representative genomes completed in 103 minutes on 16 cores with a peak memory use of 72.3 GB. All-to-all ANI estimation over the complete 5,217-genome set took <1 minute. The two dominant costs were component graph construction (58 minutes, 56.5% of wall time) and graph topology assessment (34 minutes, 33%), the former because the per-subtree progressive alignments, although independent and run in parallel, are each bounded by within-genus divergence rather than the full breadth of the family.

## Discussion

The success of PANIMaSTA in generating a single-component pangenome graph for *Anelloviridae* underscores the benefits of using general graph-theoretical techniques to inform pangenome graph construction approaches. By leveraging all-to-all pairwise genome similarity to inform maximum spanning tree (MST) to integrate globally divergent sequences via locally similar intermediates rather than base assembly directly on all-to-all pairwise alignment information, PANIMaSTA avoids the fragmentation seen in current pangenome graph construction pipelines. Notably, while a graph can be rendered single-component by forcing spurious alignments between unrelated sequences and the resultant traversal distance carries no meaningful information as every genome is nominally reachable from every other, PANIMaSTA’s topology reflects the global and local ANI similarities/genomic diversity. This makes the graph usable as a reference for read mapping, taxonomic classification, and primer design, since each of these applications depends not only on a genome being present in the reference but on its relative position within the graph space being phylogenetically meaningful.

The utility of PANIMaSTA extends beyond the anellovirus dataset presented here. Many viral and microbial families exhibit extreme genomic variation, and constructing representative pangenome graphs can pose similar computational and methodological challenges. Our MST-informed approach is especially well suited to metagenomic collections of small, and often circular, genomes of extreme sequence divergence, where the objective is to maximize the global similarity among the genomes co-embedded within a single graph and to yield a topology that reflects their global genome-to-genome similarities and phylogenetic distances. Because TANI is estimated by alignment rather than k-mer sketching, calibration is limited chiefly to the pruning threshold applied during network construction.

The immediate application enabled by a graph of this kind is similarity-aware sequence-to-graph mapping. Because traversal distance within the graph corresponds to both local and global pairwise genome similarities, the position at which a query sequence aligns is itself informative, so that a read or contig that maps to a given region of the graph can be placed relative to the genomes traversing that region, and the identity of its nearest embedded neighbors inferred directly from the locally embedded paths. For a family in which novel species continue to be described and in which the 69% identity threshold used to delineate species is applied to pairwise comparisons against a necessarily incomplete reference set, this offers an alternative to iterative pairwise search, since a single alignment against the graph situates a query within the known diversity of the family rather than against one reference at a time.

A second consequence is that node sharing within the graph tracks sequence relatedness rather than being simply an artifact of assembly. Genome pairs assigned to the same MST subtree shared node space in 22.4% of cases, at a median TANI of 0.60, compared to 0.6% of pairs assigned to different subtrees at a median TANI of 0.11. Neither extreme would be desirable, as pervasive cross-subtree sharing is expected for a tangled or hairball-like graph, in which unrelated sequences share nodes and graph position provides no phylogenetic information, while a complete absence of node sharing would prohibit cross-subtree alignment altogether. Because cross-genus node sharing is low but non-zero, the bridge genomes clearly connect disparate subtrees and can therefore enable divergent or recombinant genome sequences to similarly traverse these subtree boundaries during sequence-to-graph alignment. In a highly fragmented assembly, such sequences would be unlikely to successfully map and bridge multiple components that share no graph space. Given the importance of the Anelloviridae ORF1 region in species delineation, future efforts will focus on these loci. Finally, as the PANIMaSTA graphs can provide a substrate on which conserved loci can be identified across multiple levels, from closely-related species to intra-genus and cross-genus sequences, graph-derived measures of sequence similarity will be explored for their utility in primer design and phylogenetic analyses.

A limitation of the method arises from the possibility that excessively stringent pruning might exclude biologically relevant links, although the MST mitigates this risk by preserving the highest weighted connections for each genome. Another consideration is that longer viral or microbial genomes may require more computational resources and that the progressive alignment-based approach utilized by PANIMaSTA, VG msga, is likely worth replacing with a more recently developed alternative.

PANIMaSTA demonstrates that it is possible to construct a cohesive, single-component pangenome graph for thousands of highly divergent viral genomes by applying a MST strategy to navigate all-to-all pairwise genome similarity estimates and guide progressive assembly.

Unlike traditional alignment-based tools, this workflow efficiently links distant sequences through intermediate genomes, thereby reducing graph fragmentation and ensuring comprehensive coverage of viral diversity. The successful application to over five thousand anellovirus genomes highlights the potential of this approach to inform future studies of viral ecology, taxonomy, and clinical virome analysis. Critically, the resulting graph is not merely connected but structured, in that graph-space proximity recapitulates pairwise sequence identity across the full range of divergence within the family, so that the reference encodes phylogenetic relationships in a form directly accessible to sequence-to-graph alignment. By enabling a stable, unified reference graph whose topology is interpretable, PANIMaSTA lays the groundwork for similarity-aware mapping and classification of divergent viral sequences, for the identification of sequence space shared across genera, and more broadly emphasizes that careful integration of graph-theoretical concepts can surmount some of the most significant obstacles faced in the construction of metagenomic pangenome references.

## Author Contributions

Conceptualization: DD, SJC

Methodology: DD, SJC

Investigation: DD, FM, AJK, KP, SJC

Visualization: DD

Funding acquisition: AJK, SJC, DD

Project administration: AJK, SJC, DD

Supervision: AJK, SJC

Writing – original draft: DD, FM, SJC

Writing – review & editing: All authors

## Data Availability

Workflow code is available on GitHub: https://github.com/dduchen/panimasta

## Supplementary Figures

**Supplementary Figure S1.**
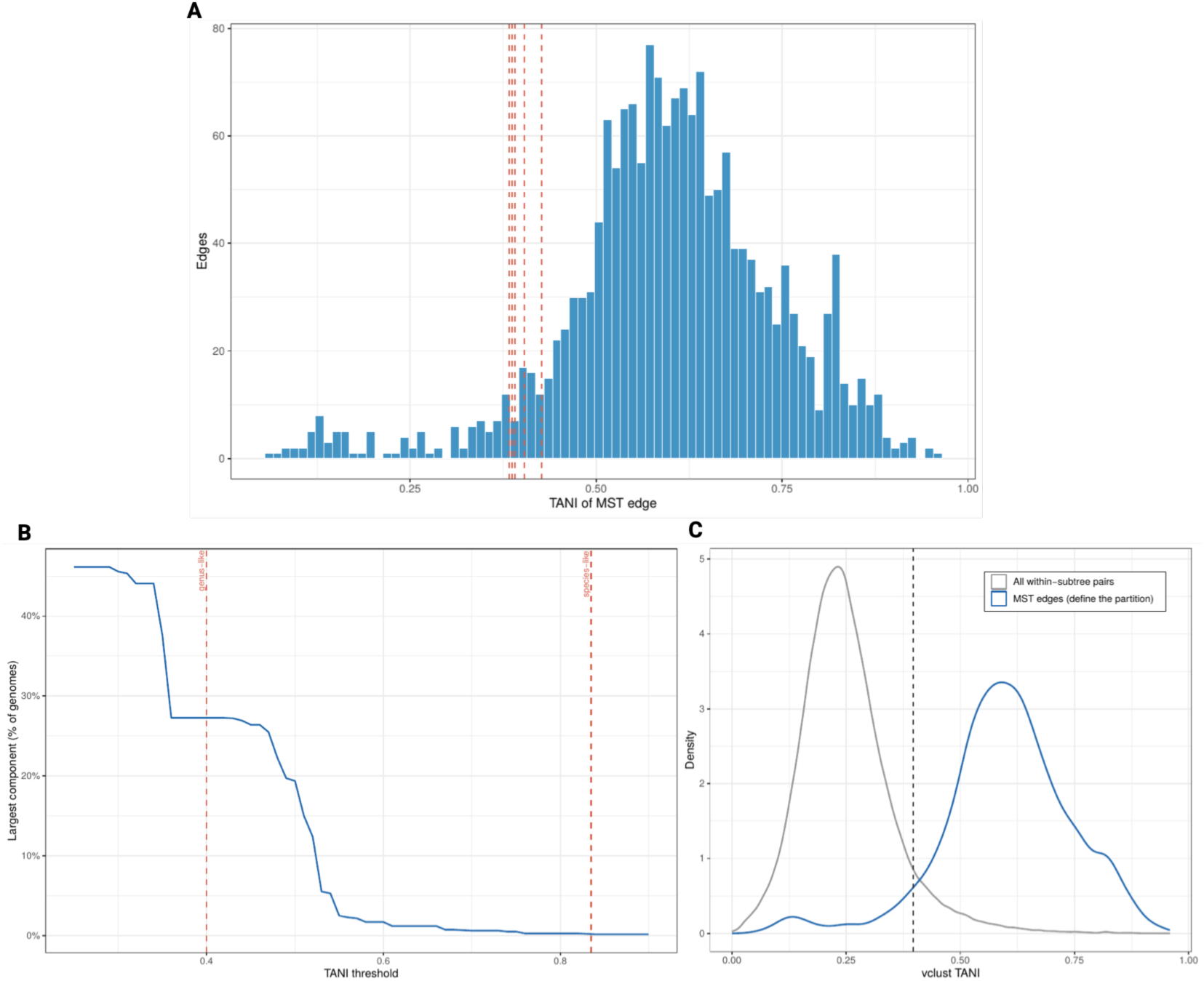
Derivation of the genus-level TANI threshold. **(A)** Distribution of TANI edge weights across MST edges. **(B)** Percolation plateau over the representative TANI matrix, with threshold of 0.397 highlighted. **(C)** Distribution of pairwise TANI estimates across all intra-subtree pairs and pairs constrained to MST edges.

**Supplementary Figure S2.**
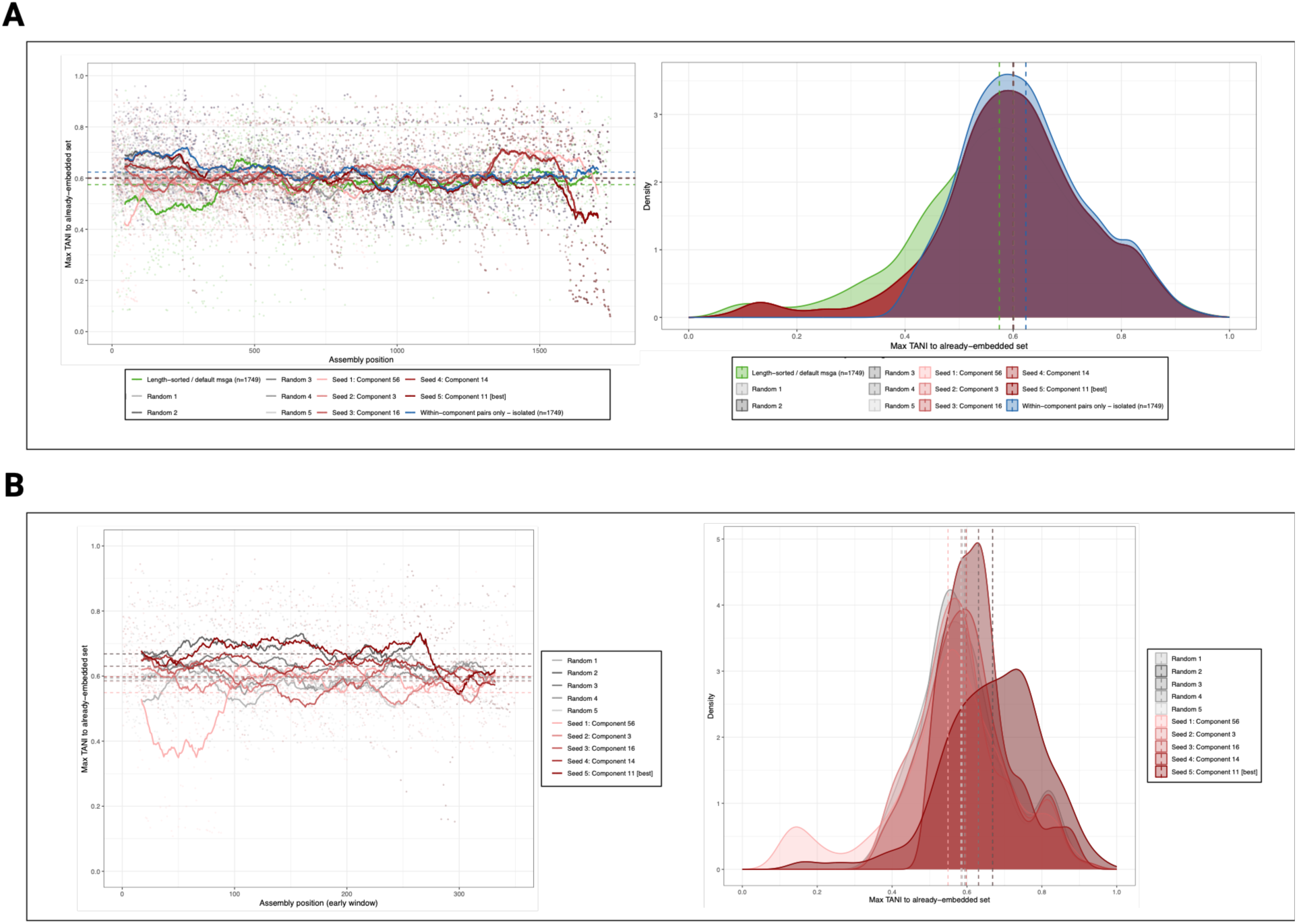
Sensitivity of the assembly to genome ordering. Progressive max TANI vs. already embedded genomes observed during assembly of the representative genomes (N=1,749). **(A)** Five component-seeded orderings compared against five random orderings, alongside the isolated and length-sorted default VG msga orderings for the full set of core/representative genomes. **(B)** Max TANI vs. already embedded genomes over the initial set of genomes (N=349, 20%). In the left panel, the positional max TANI is shown while the right panel displays the distribution. In both panels, the mean max TANI for each seed is also shown.

